# HCN domain is required for HCN channel expression and couples voltage- and cAMP-dependent gating mechanisms

**DOI:** 10.1101/2020.03.02.973560

**Authors:** Ze-Jun Wang, Ismary Blanco, Sebastien Hayoz, Tinatin I. Brelidze

## Abstract

Hyperpolarization-activated cyclic nucleotide-gated (HCN) channels are major regulators of synaptic plasticity, and rhythmic activity in the heart and brain. Opening of HCN channels requires membrane hyperpolarization and is further facilitated by intracellular cyclic nucleotides (cNMPs). In HCN channels, membrane hyperpolarization is sensed by the membrane-spanning voltage sensor domain (VSD) and the cNMP-dependent gating is mediated by the intracellular cyclic nucleotide-binding domain (CNBD) connected to the pore-forming S6 transmembrane domain via the C-linker. Previous functional analysis of HCN channels suggested a direct or allosteric coupling between the voltage- and cNMP-dependent activation mechanisms. However, the specifics of the coupling were unclear. The first cryo-EM structure of an HCN1 channel revealed that a novel structural element, dubbed HCN domain (HCND), forms a direct structural link between the VSD and C-linker/CNBD. In this study, we investigated the functional significance of the HCND. Deletion of the HCND prevented surface expression of HCN2 channels. Based on the HCN1 structure analysis, we identified R237 and G239 residues on the S2 of the VSD that form direct interactions with I135 on the HCND. Disrupting these interactions abolished HCN2 currents. We then identified three residues on the C-linker/CNBD (E478, Q382 and H559) that form direct interactions with residues R154 and S158 on the HCND. Disrupting these interactions affected both voltage- and cAMP-dependent gating of HCN2 channels. These findings indicate that the HCND is necessary for the surface expression of HCN channels, and provides a functional link between the voltage- and cAMP-dependent mechanisms of HCN channel gating.

---

Hyperpolarization-activated cyclic nucleotide-modulated (HCN) channels are peculiar members of the voltage-gated potassium channel superfamily. Unlike other potassium channels, which are depolarization-activated and highly selective for K^+^, HCN channels are activated by membrane hyperpolarization and permeate both Na^+^ and K^+^ (1,2). The opening of HCN channels is further facilitated by cyclic nucleotides (1-3). Due to these unique features HCN channels serve as pacemakers that regulate rhythmic firing of neurons and cardiomyocytes (4-6). HCN channels also have a variety of other functions, including the regulation of membrane resting potential and synaptic transmission (4). In mammals, the HCN channel family contains four isoforms (HCN1-HCN4). All four isoforms are expressed in the brain where they generate hyperpolarization-activated currents (I_h_) (1,7). Mutations in HCN1 and HCN2 channels had been linked to genetic epilepsy in humans (8-12). HCN4 channels are the predominant isoform in the heart where they generate hyperpolarization-activated inward currents known as the “funny current” (I_f_) (13,14). Genetically occurring mutations in HCN4 channels are linked to sick sinus syndrome and bradycardias in humans (15,16).

Similar to other potassium channels, HCN channels are assembled from four subunits, with each subunit containing six transmembrane segments (S1-S6). S1-S4 form a voltage-sensing domain (VSD), and S5 and S6 together with the pore-forming P-loop between them form the pore domain (PD) of the channels (4,17). In their intracellular C-terminal region HCN channels contain cyclic nucleotide-binding domain (CNBD) linked to the S6 segment with the C-linker (Fig. 1a) (17,18). The VSD is responsible for the hyperpolarization-dependent opening of the channel and the CNBD enables HCN channels to bind cyclic nucleotides and regulate membrane excitability in response to the changes in the cellular cyclic nucleotide levels (1-3). It has been suggested that the voltage-dependent and cyclic nucleotide-dependent activation mechanisms are coupled. For instance, K381E mutation in the VSD increases the effect of cAMP on HCN2 currents ∼10 fold (19). Simultaneous recording of HCN2 channel currents and changes in the fluorescence of a fluorescent cAMP analog upon binding to the channels revealed that the membrane hyperpolarization enhances the cAMP affinity of the channel activation (20). Finally, computational analysis of the voltage- and cAMP-dependent activation of HCN2 channels indicated that cAMP binding alters voltage-dependent gating of HCN2 channels (21). Despite the evidence, the molecular nature of the structural and functional coupling between the VSD and CNBD is unclear.

**Figure 1.**
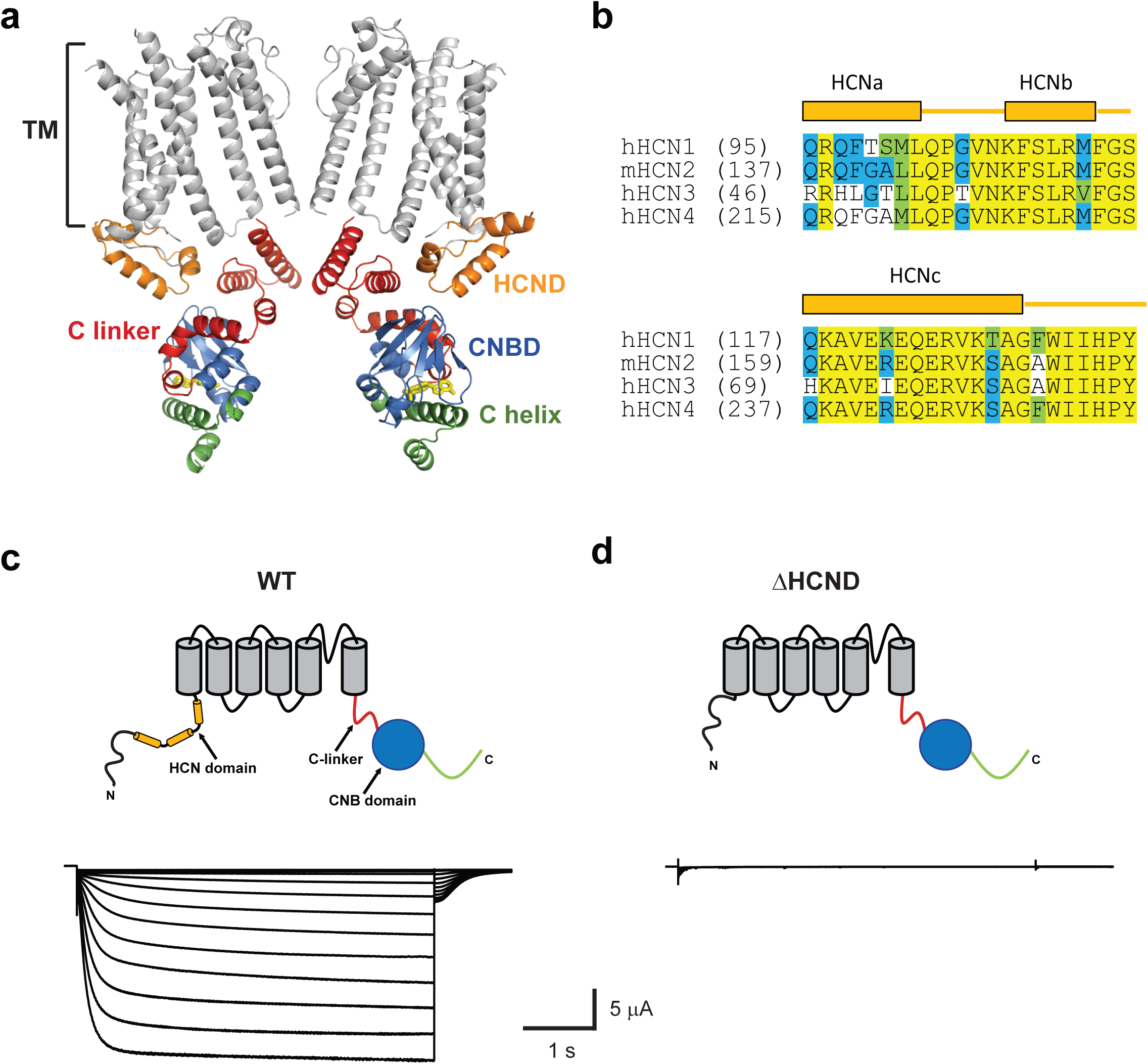
HCND is required for the function of HCN channels. **a**, Ribbon representation of the cryo-EM structure of HCN1 (pdb 5U6P (17). Only two diagonal subunits are shown for clarity. Transmembrane segments are gray, the HCN domain is orange, the C-linker is red, the CNB domain blue and the distal C-terminus green. cAMP is shown in yellow. **b**, Amino acid sequence alignment of the HCN domains for all four mammalian HCN channel isoforms. Representative currents recorded from the WT (**c**) and ΔHCND mutant (**d**) mHCN2 channels.

Recent cryo-EM structure of HCN1 channels identified a novel N-terminal structural motif formed by a stretch of 45 amino acids directly preceding the S1 transmembrane segment (Figs. 1 a and b, orange) (17). The novel motif, called HCN domain, consists of three α-helices (HCNa, HCNb and HCNc) and is conserved in all four HCN channel isoforms. Located on the periphery, the HCN domain forms direct interactions with the VSD and, also, the C-linker/CNBD (Fig. 1 a). In this study, we investigated the role of the HCN domain in the function of HCN channels by using a combination of mutagenesis, membrane surface biotinylation assay, and analysis of HCN2 channel currents recorded from excised inside-out membrane patches and with two-electrode voltage-clamp (TEVC) technique. We found that the deletion of the HCND completely abolished currents from HCN2 channels by decreasing the surface expression of the channels. Similarly, disruption of the interactions between the S2 of the VSD and HCND abolished HCN2 currents. Disruption of the interactions between the CNBD and HCND affected voltage-dependent gating, cAMP sensitivity and kinetics of deactivation of HCN2 channels. These results indicate that the HCND is necessary for the expression of functional HCN channels and it functionally links the VSD and CNBD in HCN2 channels. Preliminary results have appeared in abstract form (22).

## RESULTS

### Deletion of the HCND abolishes currents from HCN2 channels

To investigate the importance of the HCND for the function of HCN2 channels we generated ΔHCND mutant channel lacking the HCND (deletion of residues 137-180). The WT and ΔHCND mutant HCN2 channels were expressed in *Xenopus laevis* oocytes and currents were recorded using the two-electrode voltage-clamp (TEVC) technique. While the WT channels gave rise to currents with characteristic hyperpolarization-dependent activation (Fig. 1 c), ΔHCND mutant channels did not show any hyperpolarization-activated currents (Fig. 1 d). The absence of hyperpolarization-activated currents was observed in n ≥ 60 independent experiments, performed on oocytes from different batches. These results suggest that the HCN domain is necessary for the function of HCN channels.

### Deletion of the HCND dramatically decreased surface expression of HCN channels

To test if the HCN domain is necessary for the surface expression of HCN2 channels we performed surface biotinylation assay on HEK293 cells transfected with the WT and ΔHCND mutant HCN2 channels. In addition, we used untransfected HEK293 cells as a negative control. For these experiments proteins expressed at the membrane surface were biotinylated, purified from cellular lysates using neutravidin beads and then visualized using western blotting. While HEK293 cells transfected with WT HCN2 channels showed a robust surface expression of N-glycosylated and un-glycosylated (expected molecular weight ∼100 kDa) channels, as reported previously (23,24), cells transfected with the ΔHCND mutant channels only expressed unglycosylated channels at substantially lower levels than for WT HCN2 (Fig. 2A). No HCN2 channel surface expression was detected for the untransfected HEK293 cells. Staining with the α-tubulin antibody indicated that the levels of loaded protein were similar for all biotinylation experiments (Fig. 2a). Similar results were observed in n of 3 experiments, as quantified in Fig. 2b.

**Figure 2.**
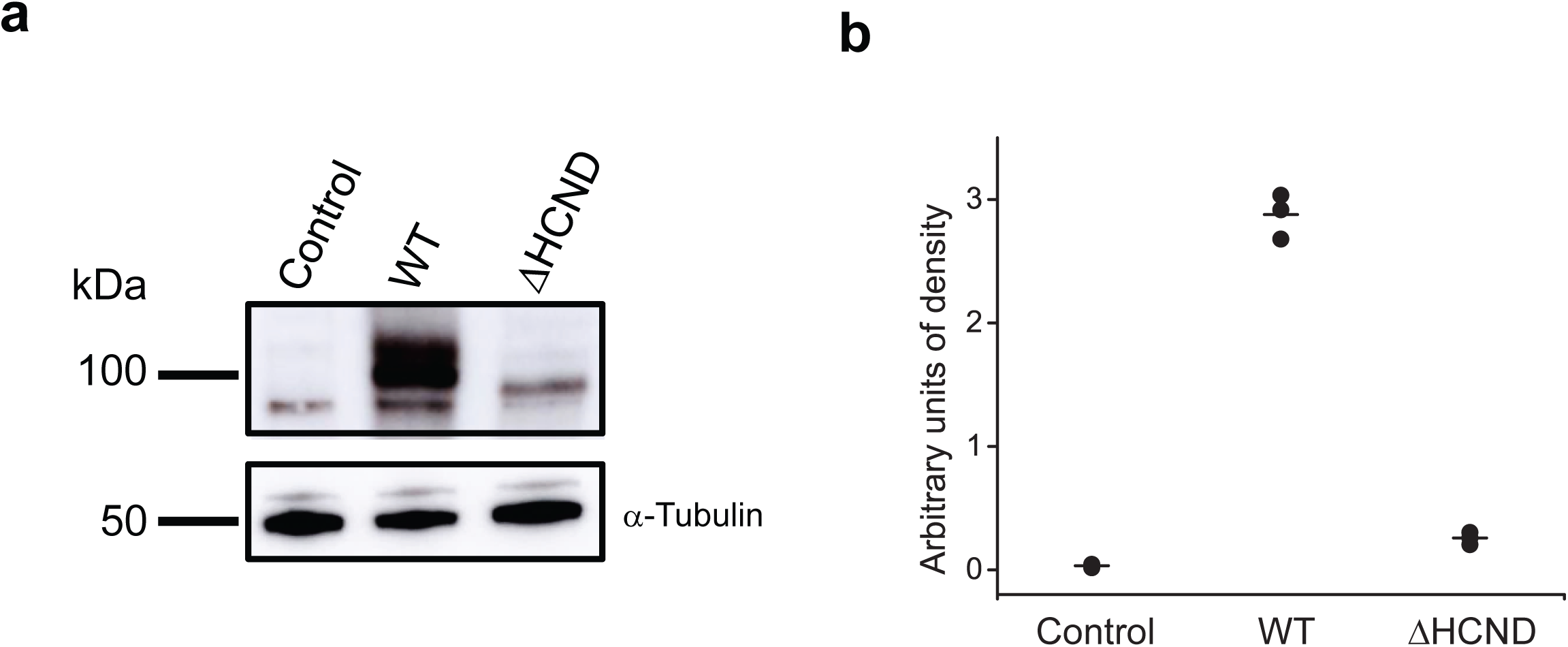
HCND is essential for the surface expression of HCN channels. **a**, Representative Immunoblot of the biotinylated protein fraction (upper half) for untransfected (control), WT and ΔHCND mHCN2 transfected HEK 293 cells probed with HCN2 specific antibodies. Representative immunoblot of the whole-cell lysates (lower half) from untransfected (control), WT and ΔHCND mHCN2 transfected HEK 293 cells probed with α-tubulin antibodies. **b**, Quantification of unglycosylated channel expression in untransfected (control), WT and ΔHCND mHCN2 transfected HEK 293 cells. n = 3.

### Neutralization of the KYK retention signal fails to rescue currents from ΔHCND channels

To investigate if deletion of the HCN domain exposed an endoplasmic reticulum (ER) retention signal we searched for subcellular localization signals in the amino acid sequence of HCN2 channels using LocSigDB, a publicly available database of experimental localization signals (25). The search identified KYK ER retention signal (26,27). The KYK ER retention signal (residues 452-454 in the mouse HCN2 channels) is located in the C-linker (Fig. 3a, left) and is conserved in all four HCN channel isoforms. Deletion of the HCN domain would increase the accessibility of the KYK residues (Fig. 3a, right). To test if neutralizing the KYK ER retention signal could rescue the currents from the ΔHCND mutant mHCN2 channels we generated mutant mHCN2 channels with alanines substituted for the KYK residues in the background of the HCND deletion (KYK-AAA/ΔHCND). Unfortunately, no hyperpolarization-activated currents were detected for the mutant channels (Fig. 3b). Therefore, neutralizing the KYK retention signal failed to rescue loss of HCN2 currents due to the HCND deletion.

**Figure 3.**
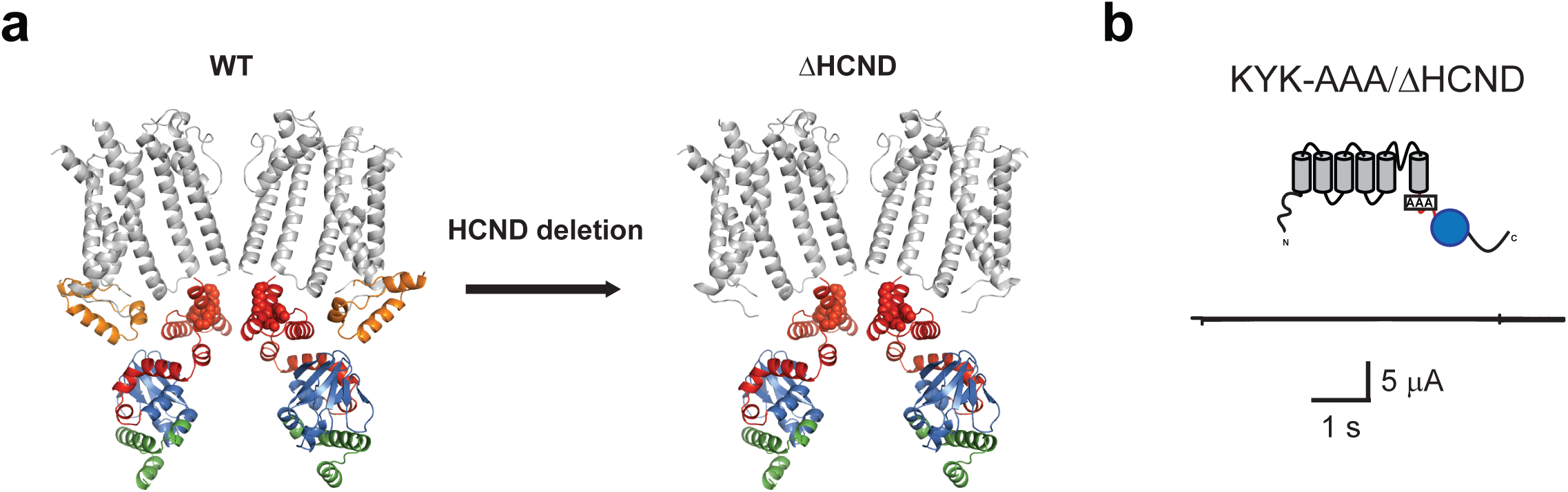
Mutating KYK ER retention signal fails to rescue ΔHCND channels. **a**, Ribbon representation of the WT and ΔHCND mutant mHCN2 channels. The same color coding as in Fig. 1a, except the KYK ER retention signal residues are shown in red spheres. **b**, Representative currents from KYK_AAA/ΔHCND mutant channels.

### Disrupting interactions between the S2 of the VSD and HCND abolishes currents from HCN channels

In addition to the importance for the targeting of HCN channels to the surface, the HCND could also contribute to the gating of HCN channels. To test the functional importance of HCND we analyzed the contacts it forms with the VSD. Analysis of the HCN1 cryo-EM structure revealed that residues R195 and G197 (R237 and G239 in HCN2) on the S2 of the VSD directly interact with the residue I135 (I177 in HCN2) on the HCND (Fig. 4a). To determine the importance of this interaction for the HCN2 channel function we mutated R237 and G239 to alanines. To our surprise, expression of G239A/R237A mutant channels in oocytes did not generate any hyperpolarization-activated currents (Fig. 4b). The absence of hyperpolarization-activated currents was observed in n ≥ 60 independent experiments, performed on oocytes from different batches. This result underscores the importance of the VSD/HCND interactions for the functional expression of HCN2 channels.

### Disrupting interactions between the CNBD and HCND affects both voltage- and cAMP-dependent gating of HCN2 channels

In the HCN1 channel cryo-EM structure, in addition to interacting with the VSD, the HCND also interacts with the CNBD. The structural analysis indicted that residues E436 and Q440 (E478 and Q482 in HCN2) on the C-linker directly interact with the residue R112 (R154 in HCN2) of the HCND of the adjacent subunit, and H517 (H559 in HCN2) in the CNBD directly interacts with the residue S116 (S158 in HCN2) in the HCND of the adjacent subunit (Fig. 5a). If the HCND forms a functional link between the VSD and CNBD then disrupting these interactions should change both the voltage-dependent and cyclic nucleotide-dependent gating of the channels. To test this hypothesis, we generated a triple mutant HCN2 channels (3M - E478A _Q482A_H559A) with the three residues on the C-linker/CNBD that interact with the HCND mutated to alanines. The 3M mutant channels gave rise to the hyperpolarization-activated currents (Fig. 5b). Analysis of the conductance versus voltage relationship indicated that the average V_1/2_ was −91.3 ± 0.3 (n = 9) for WT channels and −97.1 ± 0.3 mV (n = 10) for the 3M mutant channels (Fig. 5c). Therefore, disrupting the interactions between the CNBD and HCND shifted the voltage dependence of the channel activation to more hyperpolarized voltages by ∼ 6 mV. This is a small but statistically highly significant shift (P = 0.0001 by the unpaired Student’s t-test). The shift in the voltage-dependence suggests that the HCND functionally couples the CNBD and VSD domains in HCN2 channels.

**Figure 4.**
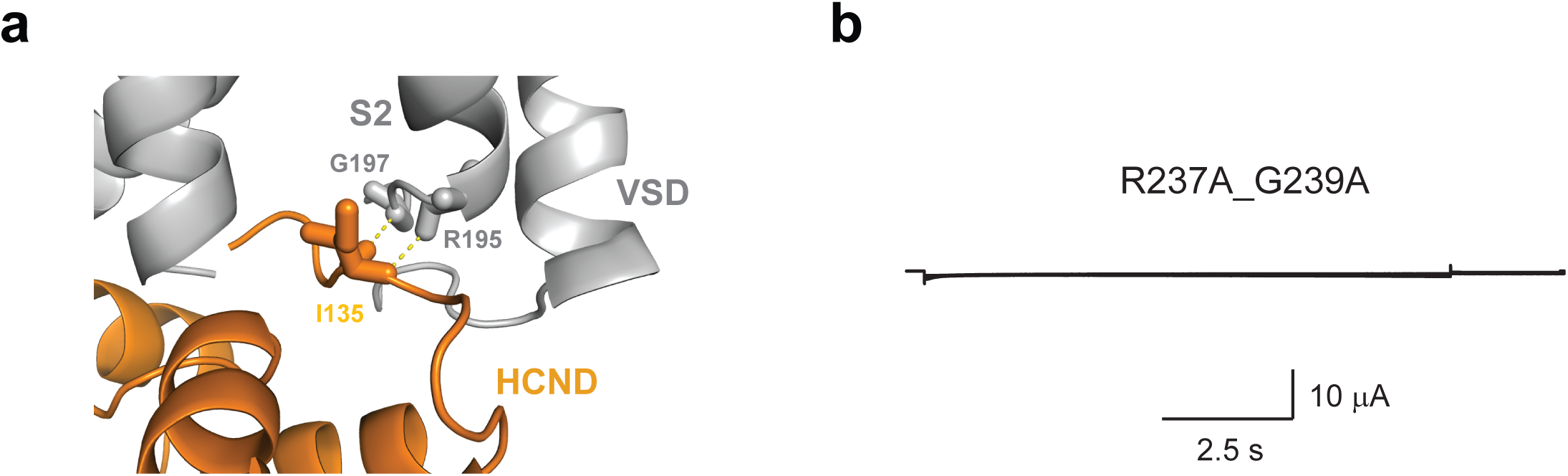
Interactions between the HCND and S2 are required for HCN channel function. **a**, Enlarged view of the interaction interface between the HCND and the VSD in HCN1 channel structure. The interacting residues G197, R195 on the S2 and I135 on the HCND are shown as sticks. **b**, Representative currents from R237A_G239A mutant mHCN2 channels. R237 and G239 correspond to R195 and G197 in HCN1 channels.

**Figure 5.**
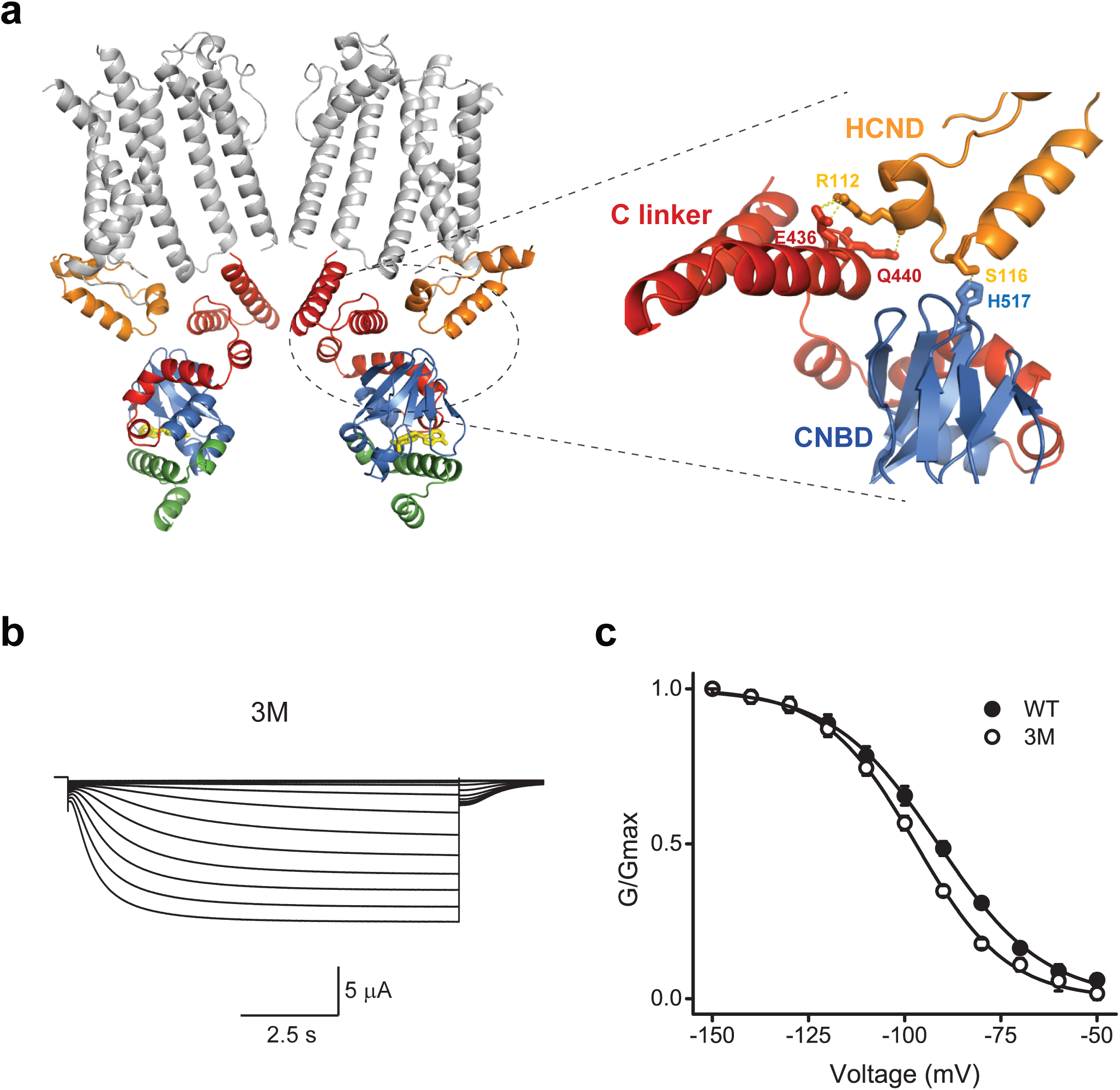
Interactions between the HCND and C-linker/CNBD affect voltage-dependent gating of HCN2 channels. **a**, Enlarged view of the interaction interface between the HCND and the C-linker/CNBD in HCN1 channels. The interacting residues E436 and Q440 on the C-linker, H517 on the CNBD and R112 on the HCND are shown as sticks. **b**, Representative currents from E478A_Q482A_H559A (3M) mutant mHCN2 channels. E478, Q482 and H559 correspond to E436, Q440 and H517 in HCN1 channels. **c**, Averaged conductance-voltage relationship for the currents from WT (black circles, n of 9) and 3M mutant (red circles, n of 10) mHCN2 channels. Lines correspond to the fits with the Boltzmann function with the V_1/2_ of −91.3 ± 0.3 mV for WT and −97.1 ± 0.3 mV for the 3M mutant, and s of 13.8 ± 0.2 and 11.9 ± 0.3 for 3M mutant mHCN2 channels.

To test if disrupting the interactions between the HCND and C-linker/CNBD also affects cyclic nucleotide-dependent activation of HCN2 channels we examined currents from WT and 3M channels over a range of cAMP concentrations. These experiments were performed using excised inside-out patch-clamp configuration, where cAMP, at the indicated concentrations, was directly applied to the intracellular side of HCN channels containing the CNBDs. Application of 10 μM cAMP increased the steady-state and tail-currents for both WT (Fig. 6a) and 3M mutant channels (Fig. 6b). The average increase was 58 ± 12 % for WT channels and 86 ± 15 % for 3M mutant channels in the presence of 10 μM cAMP (Fig. 6c). The increase in the cAMP-dependent current facilitation was greater for the 3M channels for all tested cAMP concentrations (Fig. 6c). The difference was statistically significant (P < 0.001 by two-way ANOVA), suggesting that the magnitude of the cAMP effect (potency) is increased in 3M mutant channels. The apparent cAMP affinity was 0.1 ± 0.04 μM for WT and 0.06 ± 0.02 μM for 3M mutant channels. The difference in the affinities was not statistically significant, as calculated by unpaired Student t-test.

**Figure 6.**
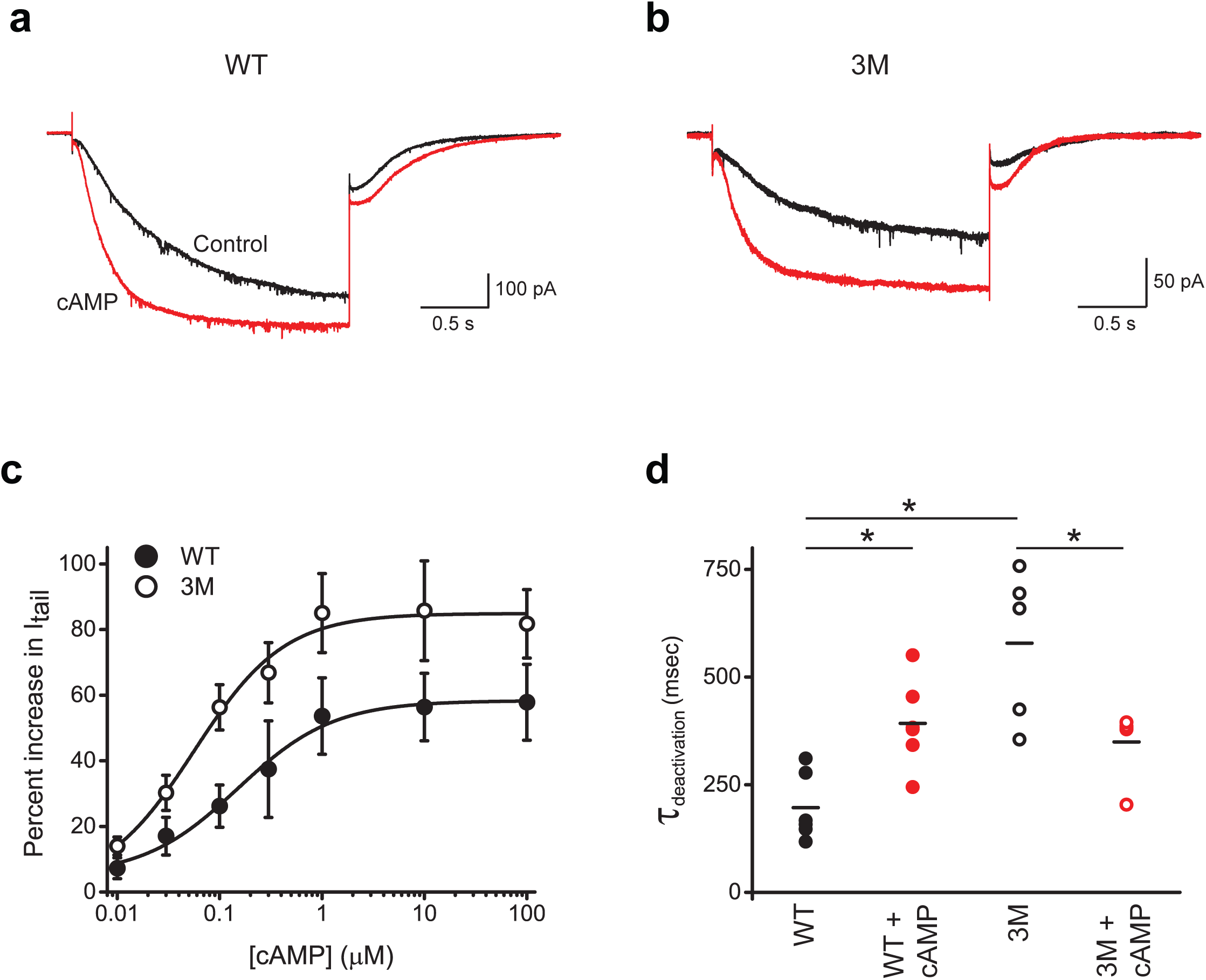
Interactions between the HCND and C-linker/CNBD affect cAMP-dependent gating of HCN2 channels. Representative currents from WT (**a**) and 3M mutant (**b**) mHCN2 channels recorded at −90 mV in the absence (black) and presence (red) of 10 μM cAMP. **c**, Plots of the percent increase in tail currents versus the cAMP concentration for WT (black circles) and 3M (red circles) mutant mHCN2 channels. The tail currents were recorded at −40 mV after the test pulse to −90 mV. Lines correspond to the fits with the Hill equation with the Kd of 0.1 ± 0.04 μM for WT and 0.06 ± 0.2 μM for 3M mutant mHCN2 channels. n ≥ 5 for each condition. **d**, Plots of deactivation time constants for tail currents recorded at −40 mV after the test pulse to −90 mV for WT and 3M mutant channels in the presence and absence of 10 μM cAMP, as indicated. n = 6 for WT and n = 5 for 3M mutant channels. The data with and without cAMP for WT and 3M channels were compared using paired Student t-test. The data between WT and 3M mutant channels were compared using unpaired Student t-test. * P < 0.02.

We also analyzed the effect of the 3M mutation on the kinetics of HCN current activation and deactivation in the absence and presence of 10 μM cAMP. The time constants of activation and deactivation were determined by fitting the activation currents and tail currents, correspondingly, with a single exponential function. While the average time constants of activation were statistically similar for WT and 3M mutant channels, both in the absence and presence of cAMP, the time constant of deactivation was larger for 3M mutant channels (Fig. 6d). This indicates that the 3M mutation slows down deactivation in the absence of cAMP. Interestingly, if for WT channels the deactivation was slower in the presence of cAMP, for 3M mutant channels cAMP accelerated the deactivation. These findings indicate that disrupting interactions between the HCND and C-linker/CNBD alters the energetics of CNBD interaction with the rest of the channel and, therefore, affects not only voltage-dependent (Fig. 5d) but also cyclic nucleotide-dependent gating.

## DISCUSSION

In this study we investigated the functional importance of the HCND and direct contacts it forms with the VSD and CNBD for HCN2 channel gating. Deletion of the HCND inhibited surface expression of HCN2 channels and disruption of the contacts between the HCND and VSD abolished currents from HCN2 channels. Therefore, the HCND is required for the expression of functional HCN channels. Disruption of the direct contacts between the HCND and C-linker/CNBD affected both voltage- and cyclic nucleotide-dependent gating of HCN channels, suggesting that the HCN domain provides not only structural but also functional link between the voltage- and cyclic nucleotide-sensing mechanisms in HCN channels.

Previous studies have shown that the amino-terminus is essential for the surface expression of HCN channels. Deletion of the entire amino-terminus (residues 1-185) abolished currents from HCN2 channels and prevented their surface expression, as indicated by the confocal images of the WT and N-terminal deleted HCN2 channels fused with GFP (28). Similarly, HCN1 channels lacking the N-terminus were retained in the ER (29). Our results show, that even if the first 136 N-terminal residues are intact but 45 residues forming the HCND (residues 137-180) are missing, expression of HCN2 channels at the surface is drastically decreased and currents are abolished (Figs. 1d and 2). Interestingly, the mutant ΔHCND HCN2 channels were still detected at the surface but at levels substantially lower than WT channels. While for the WT channels Western blot analysis showed both glycosylated and non-glycosylated populations at the surface, for ΔHCND channels no glycosylated population was detected. It has been shown that the N-glycosylation is present in native HCN channels in the brain and recombinant HCN channels expressed in HEK293 cells (24). Inhibiting N-linked glycosylation with tunicamycin and mutating the putative glycosylation site (N380) abolished currents and prevented surface expression of HCN2 channels in HEK293 cells (24). In our experiments, absence of the glycosylated ΔHCND mutant channel at the surface suggests that the deletion of the HCND prevents N-terminal glycosylation most likely by interfering with the proper folding of the protein.

Our study indicates that, in addition to affecting the surface expression, HCND is also an important functional element for fine-tuning both cNMP and voltage-dependent gating of HCN channels. HCN channel opening requires membrane hyperpolarization (1,2). Although, the dislocation of the VSD is substantial, as deduced based on the changes in the transition metal ion fluorescence resonance energy transfer (tmFRET) with membrane hyperpolarization (30) and structural analysis of the cross-linked HCN1 channels with the VSD in a hyperpolarized state (31), the movement of the VSD in response to the hyperpolarization that results in the channel opening is not clear. The VSD is connected to the pore domain (PD) via the S4-S5 linker (17). Mutations in the S4-S5 linker affect voltage-dependent gating of HCN channels (32). However, the S4-S5 linker is not required for the HCN channel opening with hyperpolarization as co-expression of the VSD and pore domain of spHCN (sea urchin HCN) channels as separate proteins generated hyperpolarization activated currents (33). It has been proposed that an extensive interface between the S4 and S5 in HCN channels is responsible for the apparent hyperpolarization-dependent activation in HCN channels (34). Consistent with this hypothesis, co-expression of the PD with the VSD lacking the C-terminal end of the S4 resulted in cNMP modulated but voltage-independent channels (33) and cross-linking the S4 and S5 could lock the channels in the “locked-open” or “locked-closed” states (35,36). More recent structural analysis of the cross-linked HCN1 channels and molecular dynamics simulations indicated that during the hyperpolarization-dependent gating the S4 helix breaks into two helices (31,37). Since the HCND forms direct interactions with the VSD, it could also contribute to the VSD movement during the voltage-dependent gating. Surprisingly, disrupting the VSD and HCND interactions with a double mutation R237A_G239A in the S2 of the VSD resulted in non-functional channels (Fig. 4b). The reason for such a dramatic effect is not clear. Possible explanations could be that the S2/HCND interactions are crucial for the function of HCN channels, and/or that the R237 and G239 residues are important for the folding of the channel.

It is thought that in the absence of cyclic nucleotides the intracellular C-linker/CNBD inhibits HCN channel opening and binding of cyclic nucleotides removes this autoinhibition (1,2). The interface between the intracellular C-linker/CNBDs from different subunits is formed by the “elbow-on-shoulder” interactions, where the elbow formed by A′-B′ alpha helices of the C-linker of one subunit is resting on the shoulder formed by the C′-D′ alpha helices of the C-linker of an adjacent subunit (18,38). In this manner, the C-linkers bring together the CNBDs in a tetrameric ring that undergoes conformational changes upon cyclic nucleotide binding, facilitating the channel opening (38-40). The HCN1 channel structure revealed that in addition to the “elbow-on-shoulder” interactions, the C-linker/CNBDs also form intersubunit interactions with the HCNDs (Fig. 5a). Therefore, cNMP-dependent gating of HCN channels most likely is accompanied not just by the rearrangement of the “elbow-on-shoulder” interface but also of the interface between the HCND and C-linker/CNBD. Our results show that breaking the intersubunit interactions between the HCND and C-linker/CNBD affects the CNBD-dependent gating of HCN2 channels by increasing the potency of HCN current facilitation by cAMP, slowing down the current deactivation in the absence of cAMP and accelerating the deactivation in the presence of cAMP (Fig. 6d). This suggests that the intersubunit interactions between the HCND and C-linker/CNBDs change the energetics of channel gating by the CNBD.

Another important conclusion of the HCN1 structure is that the HCND provides a direct structural link between the VSD and C-linker/CNBD. Here we tested the functional significance of this structural link. In addition to the effect on the cAMP-dependent gating, as discussed above, disrupting the interactions between the HCND and C-linker/CNBD domain shifted the conductance versus voltage plots to more hyperpolarized potentials (Fig. 5c). These finding suggests that the HCND provides a direct functional link between the voltage- and cNMP-dependent gating mechanisms and decreases the energetic barrier for channel activation with hyperpolarization. Consistent with our finding, Yellen and coworkers have shown that cross-lining the C-linker with the S4 in the VSD could result in “locked-open” or “lock-closed” channels (35).

At the final stages of our study Porro and coworkers published a study on the role of the HCND in HCN channel gating (41). Although, Porro and coworkers also tested effects of disrupting the interactions between the HCND and VSD and C-linker/CNBD in HCN2 channels with mutations, they considered mostly different residues in their study. For instance, we disrupted the HCND interactions with the C-linker/CNBD with a triple mutation E478A_Q482A_H559A, while Porro and coworkers examined individual and double mutations K464A and E478A. In our study, the triple mutation resulted in current facilitation by cAMP and shifted the voltage-dependence of HCN channel to more hyperpolarized potentials. For the double mutant K464A-E478A Porro and coworkers observed almost complete loss of cAMP sensitivity but no shift in the voltage-dependence of the channel activation. Our structural analysis indicates that R154, the interacting partner of E478 on the HCND, also forms interactions with Q482 on the C-linker. Therefore, the difference in the functional effect could reflect an incomplete disruption of the interactions between the HCND and C-linker. Importantly, taking together the effect of all examined mutations, Porro and coworkers concluded that the HCND is required for the proper surface expression of HCN channels and provides a functional link between the voltage- and cNMP-dependent gating mechanisms in HCN channels. It is reassuring that despite examining different interactions both our study and the study by Porro and coworkers came to the same conclusions.

Recently, it has been proposed that the voltage-dependent gating of HCN and related depolarization-activated KCNH channels is more conserved than previously thought (34). The VSD of HCN channels is capable of activating the channels with depolarization but this ability is masked by rapid inactivation. The HCND is conserved in HCN channels and is missing in KCNH channels. Therefore, while the other voltage-dependent mechanisms could be similar between the HCN and KCNH channel families, the HCND seems to provide unique means to fine tune the voltage and cNMP-dependence in HCN channels.

## EXPERIMENTAL PROCEDURES

### Electrophysiology

The cDNA encoding mouse HCN2 channels in pGEMHE vector was kindly provided by Steven Siegelbaum (Columbia University, New York, NY). Mutant HCN2 channels used in the study were generated by Bio Basic Inc. and subcloned into pGEMHE vector. The amino acid sequence of the mutant channels was confirmed using DNA sequencing (Genewiz). The cRNA was transcribed using the T7 mMessage mMachine kits (ThermoFisher Scientific). Defolliculated Xenopus *laevis* oocytes were purchased from Ecocyte Bioscience (Austin, TX), injected with the cRNA using a Nanoinject II oocyte injector (Drummond) and incubated at 18°C. HCN2 currents were recorded at room temperature with either two-electrode voltage-clamp (TEVC) amplifier (OC-725C, Warner Instruments) or with Axopatch 200A patch-clamp amplifier (Molecular Devices) for excised inside-out membrane patches. The signals were digitized using Digidata 1550 and pClamp11 software (Molecular Devices).

For TEVC experiments oocytes were placed into a RC-3Z chamber (Warner Instruments). Glass pipettes were pulled from borosilicate glass and had resistances of 0.8-1.5 MΩ when filled with 3 M KCl. The recording (bath) solution contained 100 mM KCl, 10 mM HEPES, 1 mM EGTA, pH 7.3. HCN currents were elicited by applying a series of 5 s voltage pulses (ranging from −150 to −40 mV in 10 mV increments) from a holding potential of 0 mV, followed by a 1 s voltage tail pulse to −40 mV. Currents were not leak subtracted. For currents recorded from excised membrane patches, the oocytes, following a manual removal of the vitelline membrane, were transferred to a handmade chamber containing the bath solution. Patch pipettes were pulled from borosilicate glass and had resistance of 0.5 – 1.2 MΩ after fire polishing. The extracellular (pipette) solutions contained 130 mM KCl, 10 mM HEPES, 0.2 mM EDTA, pH 7.2. The intracellular (bath) solution was either the same as the pipette solution or contained cAMP at the indicated concentrations. HCN currents were elicited by applying a 5 s voltage pulses at −90 mV from a holding potential of 0 mV, followed by a 1 s voltage tail pulse to −40 mV.

To analyze voltage dependence of HCN channel activation, peak tail-current amplitudes were normalized to the largest peak tail-current amplitude (Gmax). The normalized amplitudes were then plotted against the test voltage, and fitted with a Boltzmann equation:

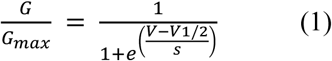

Where V represents the test voltage (mV), V_1/2_ is the midpoint activation voltage (mV), and s is the slope of the relation (mV).

cAMP was purchased from Sigma. cAMP stock was prepared in the bath solution and then diluted to obtain the range of concentrations used for the dose-response experiments. The bath solution was changed using a gravity-fed solution changer. To determine the apparent cAMP affinity (Kd), the percent increase in the peak tail current recorded after the test pulse to −90 mV was plotted versus the concentration of cAMP and fitted with a Hill equation:

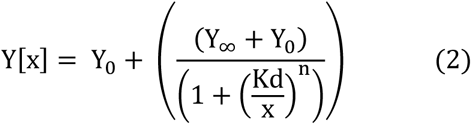

Where Y_0_ represents the minimum % increase and Y_∞_ the maximum, and n is the Hill coefficient.

The data analysis and fitting of the plots were performed in Clampfit (Molecular Devices) and Origin (Microcal Software, Inc). The error bars on the figures correspond to the SEM. Statistical analysis was performed using paired or unpaired Student’s t-tests and two-way ANOVA, as indicated. P values < 0.05 were considered significant. n represents the number of recordings from different oocytes (for TEVC experiments) or patches (for current recordings from inside-out membrane patches).

### Channel expression in HEK293 cells

For the surface biotinylation experiments, mHCN2 channels were transfected into human embryonic kidney 293 (HEK293) cells. The cDNA encoding mHCN2 channels in pcDNA3 vector for mammalian expression was kindly provided by Juliane Stieber and Andreas Ludwig (Institute for Pharmacology, Friedrich-Alexander-University, Erlangen-Nuremberg, Germany). The mutant cDNA with the deletion of the HCND was synthesized by Bio Basic Inc. and subcloned into pcDNA3 vector. The amino acid sequence of the synthesized gene was confirmed using DNA sequencing (Genewiz).

HEK293 cells were cultured and transfected with the wild-type (WT) and mutant mHCN2 channels as described in (42). 24 hrs prior to transfection, cells were plated at a cell density of 1-3×10^5^ cells into 60mm dishes. The transfection was performed using the TransIT-LT1 transfection reagent (Mirus) according to the manufacturer’s protocol.

### Biotinylation assay

All biotinylation steps were performed at 4 °C. At 48 hrs post-transfection, cells were washed twice with ice-cold phosphate buffered saline (PBS) and incubated in PBS with 1mg/mL EZ-Link Sulfo-NHS-SS-Biotin (ThermoFisher Scientific, PI21331) for 30 min. The biotin containing solution was then removed and unreacted biotin was quenched by washing the cells with 50 mM Tris-Cl, pH 7.5 for 20 minutes. The cells were then washed three times with ice-cold PBS and lysed with the buffer containing 1% Triton X-100, 150mM NaCl, 20mM EDTA, 10mM EGTA, 25mM Tris-Cl, pH 7.4 and 5 mg/ml of leupeptin, antipain, and pepstatin. The lysates were centrifuged for 15 min at 15,000 g and supernatants were transferred to a clean 1.5-ml microcentrifuge tubes containing immobilized neutravidin beads (ThermoFisher Scientific, PI29200) washed with ice-cold PBS. After 2 hrs of incubation with rotator, the tubes were centrifuged for 5 min at 2,000 g. The beads were washed five times with PBS supplemented with 0.1% SDS and centrifuged between washes.

### SDS-PAGE and Western Blotting

The biotinylated proteins were eluted off the beads by adding 2x Laemmli Sample Buffer at 95 °C for 5 min and loaded on 12% Bis-Tris SDS-polyacrylamide gels. After SDS-PAGE, proteins were transferred to a nitrocellulose membrane by electroblotting. The HCN channels were detected using a rabbit anti-HCN2 primary antibody (Boster, A02804) that recognizes the C-terminal sequence of HCN2 channels and a goat anti-rabbit secondary antibody (Abcam, ab6721). Chemiluminescent signal was detected using SuperSignal West Femto Maximum Sensitivity Substrate Chemiluminescence detection kit (Pierce). As a loading control we used α-tubulin antibody (ThermoFisher Scientific, NB100690SS). The band intensities were analyzed with ImageJ.

## Acknowledgments

We are grateful to William N Zagotta for helpful discussions. We would like to thank Gerard Ahern for generously allowing us to use his lab equipment to carry out the initial experiments for the project, and to Robert Yasuda for providing the HEK293 cell cultures for our experiments. This work was supported by the National Institute of General Medicine grant R01GM124020 (T.I.B.).

## Authorship contributions

T.I.B. conceived the study. Z.J.W, I.B. and S.H. performed the experiments. Z.J.W, I.B. S.H. and T.I.B. performed data analysis. T.I.B. wrote the manuscript with the input from all coauthors.

## Materials & correspondence

Tinatin I. Brelidze

## REFERENCIES

1. Ludwig, A., Zong, X., Jeglitsch, M., Hofmann, F., and Biel, M. (1998) A family of hyperpolarization-activated mammalian cation channels. Nature 393, 587–591

2. Santoro, B., Liu, D. T., Yao, H., Bartsch, D., Kandel, E. R., Siegelbaum, S. A., and Tibbs, G. R. (1998) Identification of a gene encoding a hyperpolarization-activated pacemaker channel of brain. Cell 93, 717–729

3. DiFrancesco, D., and Tortora, P. (1991) Direct activation of cardiac pacemaker channels by intracellular cyclic AMP. Nature 351, 145–147

4. Wahl-Schott, C., and Biel, M. (2009) HCN channels: structure, cellular regulation and physiological function. Cell Mol Life Sci 66, 470–494

5. Kaupp, U. B., and Seifert, R. (2001) Molecular diversity of pacemaker ion channels. Annu Rev Physiol 63, 235–257

6. DiFrancesco, D. (2010) The role of the funny current in pacemaker activity. Circ Res 106, 434–446

7. Santoro, B., Chen, S., Luthi, A., Pavlidis, P., Shumyatsky, G. P., Tibbs, G. R., and Siegelbaum, S. A. (2000) Molecular and functional heterogeneity of hyperpolarization-activated pacemaker channels in the mouse CNS. J. Neurosci 20, 5264–5275

8. Dibbens, L. M., Reid, C. A., Hodgson, B., Thomas, E. A., Phillips, A. M., Gazina, E., Cromer, B. A., Clarke, A. L., Baram, T. Z., Scheffer, I. E., Berkovic, S. F., and Petrou, S. (2010) Augmented currents of an HCN2 variant in patients with febrile seizure syndromes. Ann. Neurol 67, 542–546

9. Tang, B., Sander, T., Craven, K. B., Hempelmann, A., and Escayg, A. (2008) Mutation analysis of the hyperpolarization-activated cyclic nucleotide-gated channels HCN1 and HCN2 in idiopathic generalized epilepsy. Neurobiol Dis 29, 59–70

10. DiFrancesco, J. C., Barbuti, A., Milanesi, R., Coco, S., Bucchi, A., Bottelli, G., Ferrarese, C., Franceschetti, S., Terragni, B., Baruscotti, M., and DiFrancesco, D. (2011) Recessive loss-of-function mutation in the pacemaker HCN2 channel causing increased neuronal excitability in a patient with idiopathic generalized epilepsy. J. Neurosci 31, 17327–17337

11. Nakamura, Y., Shi, X., Numata, T., Mori, Y., Inoue, R., Lossin, C., Baram, T. Z., and Hirose, S. (2013) Novel HCN2 mutation contributes to febrile seizures by shifting the channel’s kinetics in a temperature-dependent manner. PLoS. One 8, e80376

12. Nava, C., Dalle, C., Rastetter, A., Striano, P., de Kovel, C. G., Nabbout, R., Cances, C., Ville, D., Brilstra, E. H., Gobbi, G., Raffo, E., Bouteiller, D., Marie, Y., Trouillard, O., Robbiano, A., Keren, B., Agher, D., Roze, E., Lesage, S., Nicolas, A., Brice, A., Baulac, M., Vogt, C., El, H. N., Schneider, E., Suls, A., Weckhuysen, S., Gormley, P., Lehesjoki, A. E., De, J. P., Helbig, I., Baulac, S., Zara, F., Koeleman, B. P., Haaf, T., LeGuern, E., and Depienne, C. (2014) De novo mutations in HCN1 cause early infantile epileptic encephalopathy. Nat. Genet 46, 640–645

13. DiFrancesco, D. (1993) Pacemaker mechanisms in cardiac tissue. Annu. Rev. Physiol 55, 455–472

14. Shi, W., Wymore, R., Yu, H., Wu, J., Wymore, R. T., Pan, Z., Robinson, R. B., Dixon, J. E., McKinnon, D., and Cohen, I. S. (1999) Distribution and prevalence of hyperpolarization-activated cation channel (HCN) mRNA expression in cardiac tissues. Circ. Res 85, e1–e6

15. Moller, M., Silbernagel, N., Wrobel, E., Stallmayer, B., Amedonu, E., Rinne, S., Peischard, S., Meuth, S. G., Wunsch, B., Strutz-Seebohm, N., Decher, N., Schulze-Bahr, E., and Seebohm, G. (2018) In Vitro Analyses of Novel HCN4 Gene Mutations. Cell Physiol Biochem 49, 1197–1207

16. Schulze-Bahr, E., Neu, A., Friederich, P., Kaupp, U. B., Breithardt, G., Pongs, O., and Isbrandt, D. (2003) Pacemaker channel dysfunction in a patient with sinus node disease. J Clin Invest 111, 1537–1545

17. Lee, C. H., and MacKinnon, R. (2017) Structures of the Human HCN1 Hyperpolarization-Activated Channel. Cell 168, 111–120

18. Zagotta, W. N., Olivier, N. B., Black, K. D., Young, E. C., Olson, R., and Gouaux, E. (2003) Structural basis for modulation and agonist specificity of HCN pacemaker channels. Nature 425, 200–205

19. Wicks, N. L., Chan, K. S., Madden, Z., Santoro, B., and Young, E. C. (2009) Sensitivity of HCN channel deactivation to cAMP is amplified by an S4 mutation combined with activation mode shift. Pflugers Arch 458, 877–889

20. Kusch, J., Biskup, C., Thon, S., Schulz, E., Nache, V., Zimmer, T., Schwede, F., and Benndorf, K. (2010) Interdependence of receptor activation and ligand binding in HCN2 pacemaker channels. Neuron 67, 75–85

21. Hummert, S., Thon, S., Eick, T., Schmauder, R., Schulz, E., and Benndorf, K. (2018) Activation gating in HCN2 channels. PLoS Comput Biol 14, e1006045

22. Wang, Z. J., Hayoz, S., and Brelidze, T. I. (2019) The Role of HCN Domain in the Function of HCN Channels. Biophysical Journal 116, 103a–103a

23. Akhavan, A., Atanasiu, R., Noguchi, T., Han, W., Holder, N., and Shrier, A. (2005) Identification of the cyclic-nucleotide-binding domain as a conserved determinant of ion-channel cell-surface localization. J. Cell Sci 118, 2803–2812

24. Much, B., Wahl-Schott, C., Zong, X., Schneider, A., Baumann, L., Moosmang, S., Ludwig, A., and Biel, M. (2003) Role of subunit heteromerization and N-linked glycosylation in the formation of functional hyperpolarization-activated cyclic nucleotide-gated channels. J. Biol. Chem 278, 43781–43786

25. King, B. R., and Guda, C. (2007) ngLOC: an n-gram-based Bayesian method for estimating the subcellular proteomes of eukaryotes. Genome Biol 8, R68

26. Jackson, M. R., Nilsson, T., and Peterson, P. A. (1990) Identification of a consensus motif for retention of transmembrane proteins in the endoplasmic reticulum. EMBO J 9, 3153–3162

27. Nilsson, T., Jackson, M., and Peterson, P. A. (1989) Short cytoplasmic sequences serve as retention signals for transmembrane proteins in the endoplasmic reticulum. Cell 58, 707–718

28. Proenza, C., Tran, N., Angoli, D., Zahynacz, K., Balcar, P., and Accili, E. A. (2002) Different roles for the cyclic nucleotide binding domain and amino terminus in assembly and expression of hyperpolarization-activated, cyclic nucleotide-gated channels. J Biol Chem 277, 29634–29642

29. Pan, Y., Laird, J. G., Yamaguchi, D. M., and Baker, S. A. (2015) An N-Terminal ER Export Signal Facilitates the Plasma Membrane Targeting of HCN1 Channels in Photoreceptors. Invest Ophthalmol Vis Sci 56, 3514–3521

30. Dai, G., Aman, T. K., DiMaio, F., and Zagotta, W. N. (2019) The HCN channel voltage sensor undergoes a large downward motion during hyperpolarization. Nat Struct Mol Biol 26, 686–694

31. Lee, C. H., and MacKinnon, R. (2019) Voltage Sensor Movements during Hyperpolarization in the HCN Channel. Cell 179, 1582–1589 e1587

32. Chen, J., Mitcheson, J. S., Tristani-Firouzi, M., Lin, M., and Sanguinetti, M. C. (2001) The S4-S5 linker couples voltage sensing and activation of pacemaker channels. Proc Natl Acad Sci U S A 98, 11277–11282

33. Flynn, G. E., and Zagotta, W. N. (2018) Insights into the molecular mechanism for hyperpolarization-dependent activation of HCN channels. Proc Natl Acad Sci U S A 115, E8086–E8095

34. Cowgill, J., Klenchin, V. A., Alvarez-Baron, C., Tewari, D., Blair, A., and Chanda, B. (2019) Bipolar switching by HCN voltage sensor underlies hyperpolarization activation. Proc Natl Acad Sci U S A 116, 670–678

35. Kwan, D. C., Prole, D. L., and Yellen, G. (2012) Structural changes during HCN channel gating defined by high affinity metal bridges. J Gen Physiol 140, 279–291

36. Shin, K. S., Rothberg, B. S., and Yellen, G. (2001) Blocker state dependence and trapping in hyperpolarization-activated cation channels: evidence for an intracellular activation gate. J Gen Physiol 117, 91–101

37. Kasimova, M. A., Tewari, D., Cowgill, J. B., Ursuleaz, W. C., Lin, J. L., Delemotte, L., and Chanda, B. (2019) Helix breaking transition in the S4 of HCN channel is critical for hyperpolarization-dependent gating. Elife 8

38. Craven, K. B., and Zagotta, W. N. (2004) Salt bridges and gating in the COOH-terminal region of HCN2 and CNGA1 channels. J. Gen. Physiol 124, 663–677

39. Puljung, M. C., DeBerg, H. A., Zagotta, W. N., and Stoll, S. (2014) Double electron-electron resonance reveals cAMP-induced conformational change in HCN channels. Proc Natl Acad Sci U S A 111, 9816–9821

40. Wainger, B. J., DeGennaro, M., Santoro, B., Siegelbaum, S. A., and Tibbs, G. R. (2001) Molecular mechanism of cAMP modulation of HCN pacemaker channels. Nature 411, 805–810

41. Porro, A., Saponaro, A., Gasparri, F., Bauer, D., Gross, C., Pisoni, M., Abbandonato, G., Hamacher, K., Santoro, B., Thiel, G., and Moroni, A. (2019) The HCN domain couples voltage gating and cAMP response in hyperpolarization-activated cyclic nucleotide-gated channels. Elife 8

42. Gianulis, E. C., Liu, Q., and Trudeau, M. C. (2013) Direct interaction of eag domains and cyclic nucleotide-binding homology domains regulate deactivation gating in hERG channels. J. Gen. Physiol 142, 351–366

